# A Three-dimensional Analytical Framework for Retinal Microvasculature Reveals Layer-associated Vulnerability in Development and Neovascular Remodeling

**DOI:** 10.64898/2026.03.16.711909

**Authors:** Wenhao Shang, Guiying Hong, William E. Keller, Ryan A. Morton, Pierre Zeboulon, Toma Kenichi, Xin Duan, Douglas B. Gould, Tyson N. Kim

## Abstract

The neurosensory retina is one of the most metabolically active tissues in the body and a uniquely accessible extension of the central nervous system, where neuronal and vascular structures can be visualized non-invasively. Its accessibility and highly organized laminar architecture make it a powerful model for studying vascular development and a window into systemic health. Although computational analyses of retinal images have enabled risk assessment for ocular and systemic diseases, most vascular studies rely on two-dimensional frameworks with limited resolution of capillary structure and layer-specific organization. Here, we present a high-resolution three-dimensional (3D) imaging and analysis pipeline enabling quantification of retinal microvasculature and extraction of structural and network metrics across vascular layers. We apply this approach to two mouse models of aberrant retinal vascular development: one with spontaneous postnatal chorioretinal neovascularization and another with disrupted neurovascular lattice formation and layered organization in early life. Across both pathologic contexts, 3D analysis enables detailed characterization of retinal vascular architecture and identifies early vulnerability within the intermediate plexus, the vascular network that bridges the superficial and deep retinal layers, as a sensitive indicator of abnormal remodeling and neovascularization. This framework enables precise characterization of retinal vasculature and establishes a foundation for identifying new retinal biomarkers with potential relevance to neurovascular and systemic disease.

## Introduction

The retina provides a unique opportunity to study the central nervous system *in vivo*. As a direct extension of the brain, it enables non-invasive, high-resolution visualization of neural and vascular networks in their native physiological context. The mammalian retina is among the most metabolically active tissues in the body, containing hierarchical vascular plexuses arranged in parallel planes and interconnected by orthogonal penetrating vessels.^1^ This combination of structure and imaging accessibility has made the retina both a benchmark model for vascular development and a powerful window into systemic health. A growing body of evidence demonstrates that retinal vascular abnormalities provide quantitative biomarkers in identifying and monitoring cardiovascular and neurological diseases.^2,3^ Advances in computational and deep learning approaches have revealed that retinal vascular features captured in fundus images can predict systemic disease risk with accuracy comparable to clinical assessment.^4^ Further work has identified markers of cerebral small vessel diseases within the deeper retinal plexus,^5^ highlighting the relevance of pathologic features amid specific retinal layers. As imaging and analytical methods advance, a deeper understanding of the retina’s three-dimensional (3D) microarchitecture is poised to illuminate fundamental links between ocular, cerebral, and systemic health.^6^

Most retinal vascular analyses rely on two-dimensional (2D) approaches without depth resolution and primarily focus on macrovascular structures using conventional imaging such as fundus photography.^7^ By projecting all retinal layers onto a single plane and emphasizing larger vessels, these analyses can obscure microvascular alterations specific to the retina’s layers, each with distinct functional and metabolic demands. Optical coherence tomography angiography (OCTA) has emerged as a powerful tool for non-invasive, depth-resolved visualization of the retinal vasculature, enabling 3D mapping of the superficial and deep plexuses.^8,9^ Ongoing advances in OCTA-based reconstruction and quantification have begun to capture finer vascular detail, though current analyses remain largely focused on vessel-level metrics with limited representations of the intermediate plexus and inter-layer connectivity.^10–13^ Complementary advances in high-resolution optical techniques, particularly multiphoton microscopy, extend this capability by achieving *in vivo* visualization of structural and dynamic biological processes such as neovascularization, perfusion, and blood-flow dynamics within the retina with subcellular resolution,^14–16^ serving as an imaging benchmark for high-resolution analysis in animal models. Multiphoton microscopy achieves this by simultaneous absorption of two or more lower-energy photons, typically in the near-infrared range. The use of longer excitation wavelengths reduces light scattering and phototoxicity, while the nonlinear dependence of fluorescence on excitation intensity confines signal generation to the focal volume. These properties confer intrinsic optical sectioning in 3D and allow imaging at subcellular resolution to depths not accessible by conventional microscopies.^17–19^ A growing number of studies demonstrate the importance of high-resolution 3D imaging for capturing fine architectural features in the retina,^20,21^ yet quantitative frameworks to characterize these complex 3D networks and architectures remain limited.^22–24^ While computational tools for automated vascular network quantification have been developed, including 2D frameworks such as AngioTool^25^ and recent 3D approaches for deep learning-based segmentation and morphometric extraction^26^ or graph-based analysis of pre-segmented vasculature,^27^ these methods have primarily focused on vessel-level morphometrics (density, length, radius) and additional work is needed to address layer-resolved architecture, inter-plexus connectivity, or fractal complexity within individual vascular compartments. Advancing such 3D analytical methods will be valuable for capturing the retina’s layer-specific organization, inter-plexus connectivity, and topological complexity that provides deeper insight into vascular development, disease mechanisms, and potential biomarkers of retinal and systemic disorders.

The retinal vasculature comprises three distinct and highly organized plexus layers that precisely regulate blood flow and nutrient delivery to meet the retina’s high metabolic demands.^28,29^ Each layer exhibits unique structural and functional characteristics, as well as vulnerability across disease states. Despite this recognized layer-specific vulnerability, quantitative characterization of the intermediate capillary plexus has been limited, in part because most imaging and analytical frameworks struggle to resolve this compartment as an independent entity.^3,30^ Quantifying structural changes within and between these layers may yield nuanced insights into vascular development and disease pathology. Our recent work with Toma et al. demonstrated that the development of these inter-plexus vessels is regulated by a subpopulation of perivascular retinal ganglion cells (RGCs) through direct neuron-vessel contact.^31^ Prior studies have shown that these vertical vessels also arise predominantly from veins and capillaries, underscoring their distinct developmental origin.^32^ To systematically investigate such complex spatial and functional relationships, we developed a comprehensive 3D quantification framework that extracts detailed structural metrics from high-resolution imaging data, enabling quantitative analysis of retinal vascular organization across multiple layers and their interconnectivity. This framework has provided new insight into neurovascular patterning and dysgenesis, including the discovery of a novel mechanism of vascular development in the retina whereby specialized neurons guide 3D vascular lattice organization mediated through direct neuron-vessel contact.^31^ Here, we extend this framework to resolve distinct mechanisms of 3D vascular remodeling in the retina, applying it in two representative contexts relevant to development and disease: (i) spontaneous chorioretinal neovascularization in postnatal life and (ii) disrupted 3D vascular lattice formation in the developing retina.

## Results

### A pipeline for 3D retinal vasculature analysis

We developed an end-to-end pipeline for analyzing 3D retinal vasculature, which reconstructs full-thickness networks from high-resolution image Z-stacks through computer-aided segmentation, yielding detailed computational representations of the 3D vascular architecture (Fig. 1a–c). The reconstructed networks are subsequently stratified into anatomically distinct layers: the superficial layer plexus (SL), middle layer plexus (ML), and deep layer plexus(DL), along with identification of their inter-plexus connections (Fig. 1c,d).

**Figure 1.**
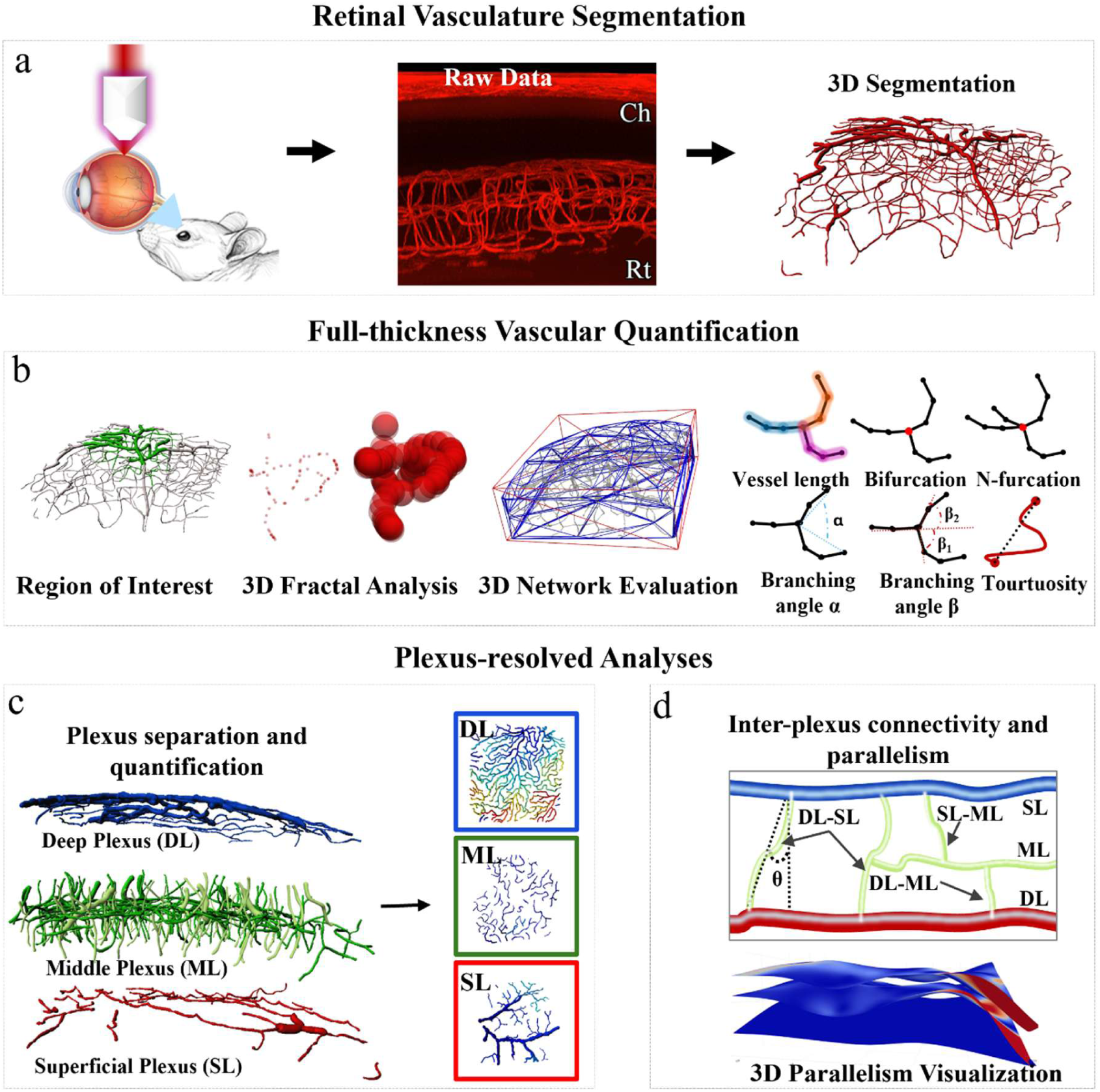
Pipeline for 3D retinal vasculature analysis. **(a)** Retinal vasculature segmentation. High-resolution two-photon excitation microscopy Z-stacks spanning the choroid (Ch) and the neurosensory retina (Rt) were acquired. The volumetric data were segmented to generate three-dimensional representations for analysis. **(b)** Full-thickness vascular quantification. Global analyses of vessel morphology and network architecture were performed across the entire vascular volume or within user-defined regions of interest. Quantified parameters include vessel length, diameter, branching angle, tortuosity, vessel volume density, branching topology, network connectivity, and fractal dimension. **(c)** Plexus-resolved analyses. The retinal vasculature was separated into the superficial layer (SL), middle layer (ML), and deep layer (DL) vascular plexuses. Plexus-specific quantification was then performed to characterize layer-dependent vascular architecture. **(d)** Inter-plexus connectivity and parallelism. Cross-plexus relationships were analyzed by characterizing direct inter-plexus connections between SL, ML, and DL and assessing deviation in inter-plexus parallelism.

Following reconstruction and layer stratification, we established a multi-level quantification framework that systematically characterizes retinal vascular metrics across different organizational scales. Our approach integrates three complementary analytical modules: (i) full-thickness analysis capturing global network properties, (ii) plexus-specific quantification revealing layer-dependent features, and (iii) inter-layer connectivity analysis characterizing cross-plexus organization, a dimension largely unexplored in previous studies. At the full-thickness and individual plexus levels, we quantify standard morphometric and topological parameters including vessel segment length, diameter, tortuosity, vessel volume density, vessel number density, and branching point density (Fig. 1b-c). In addition to these conventional metrics, we incorporate 3D fractal dimension (3D FD) to capture geometric complexity and introduce layer-specific parameters, such as the fragmentation index, to assess network characteristics.

For inter-layer connectivity analysis, we developed metrics specifically designed to quantify cross-plexus organization: (i) inter-layer parallelism, measuring the degree of parallel alignment between vascular plexuses; (ii) inter-plexus connection classification, categorizing and quantifying different connection types between specific vascular layers (DL–ML, SL–ML, DL–SL); and (iii) inter-plexus vessel geometries, characterized by excursion ratio (vessel path length normalized to vertical span) and orientation angle θ relative to the retinal plane (Fig. 1d). These metrics enable systematic detection of alterations in vertical vessel architecture and inter-layer coordination. All analyses can be performed across entire vascular territories or within user-defined regions of interest (ROI), providing flexible spatial resolution for both global assessments and targeted investigation within specific vascular domains. A comprehensive catalog of quantification parameters and their definitions is provided in Table 1. The description of the analytical workflow, including image acquisition through 3D reconstruction to multi-level quantification, is illustrated in Fig 1.

### 3D structural analysis of spontaneous chorioretinal neovascularization and anastomoses reveals capillary rarefaction, ML fragmentation, and radialization of DL vessels

We evaluated an established model of chorioretinal neovascularization, wherein mice harboring a missense point mutation of type-IV collagen (Col4a1^+/Δex41^) develop neovascularization and chorioretinal anastomoses (CRA) in postnatal life. These mice represent a robust genetic model of spontaneous chorioretinal neovascularization,^33^ providing a platform for interrogating the 3D structural features of pathologic angiogenesis in the adult eye.

3D image data of the chorioretinal vasculature were acquired from three experimental groups: littermate controls, mutant mice not exhibiting neovascular lesions (Col4a1^+/Δex41^), and mutant mice harboring neovascular changes and CRA (Col4a1^+/Δex41^+CRA). Following vascular segmentation of raw image data, we performed full-thickness network analysis and quantification of vascular characteristics within a standardized region of interest (ROI; 285 × 285 μm, matching the mean CRA lesion size; Fig. 2a, c) spanning the complete retinal vascular thickness from choroid to superficial layer plexus, ensuring consistent spatial sampling across eyes for downstream quantitative analyses. Geometric characterization at the full-thickness level revealed profound vascular remodeling in the Col4a1^+/Δex41^+CRA group. Vessels exhibited significantly larger mean diameters (CRA: 7.47 ± 3.31 μm vs. control: 3.77 ± 1.74 μm; *p*<0.0001; Fig. 2d) and increased vessel volume density (Fig. 2e) compared with both control and Col4a1^+/Δex41^ groups. In contrast, vascular network connectivity was markedly compromised, with vessel number density reduced by 72% (CRA: 4.88 ± 2.62 vs. control: 17.41 ± 10.79 ×10^-4^/μm³; *p*<0.05; Fig. 2f) and similar reductions in branching point density (Fig. 2g). To quantify microvascular architectural complexity, we computed the 3D fractal dimension (3D FD) of vascular surfaces using the Bouligand–Minkowski method (see Methods).^34^ The Col4a1^+/Δex41^+CRA group exhibited significantly elevated 3D FD (1.86 ± 0.08) compared with control (1.68 ± 0.09) and Col4a1^+/Δex41^ mice (1.67 ± 0.06), representing an ∼11% increase (*p*<0.05 for both comparisons; Fig. 2h). Additional morphometric metrics, including vessel segment count, branching points, total vessel length, mean segment length, and vessel straightness are provided in Supplementary Fig. S1. This elevation reflects greater surface irregularity and volumetric complexity, which we find to be characteristic of pathological neovascular structures.

**Figure 2.**
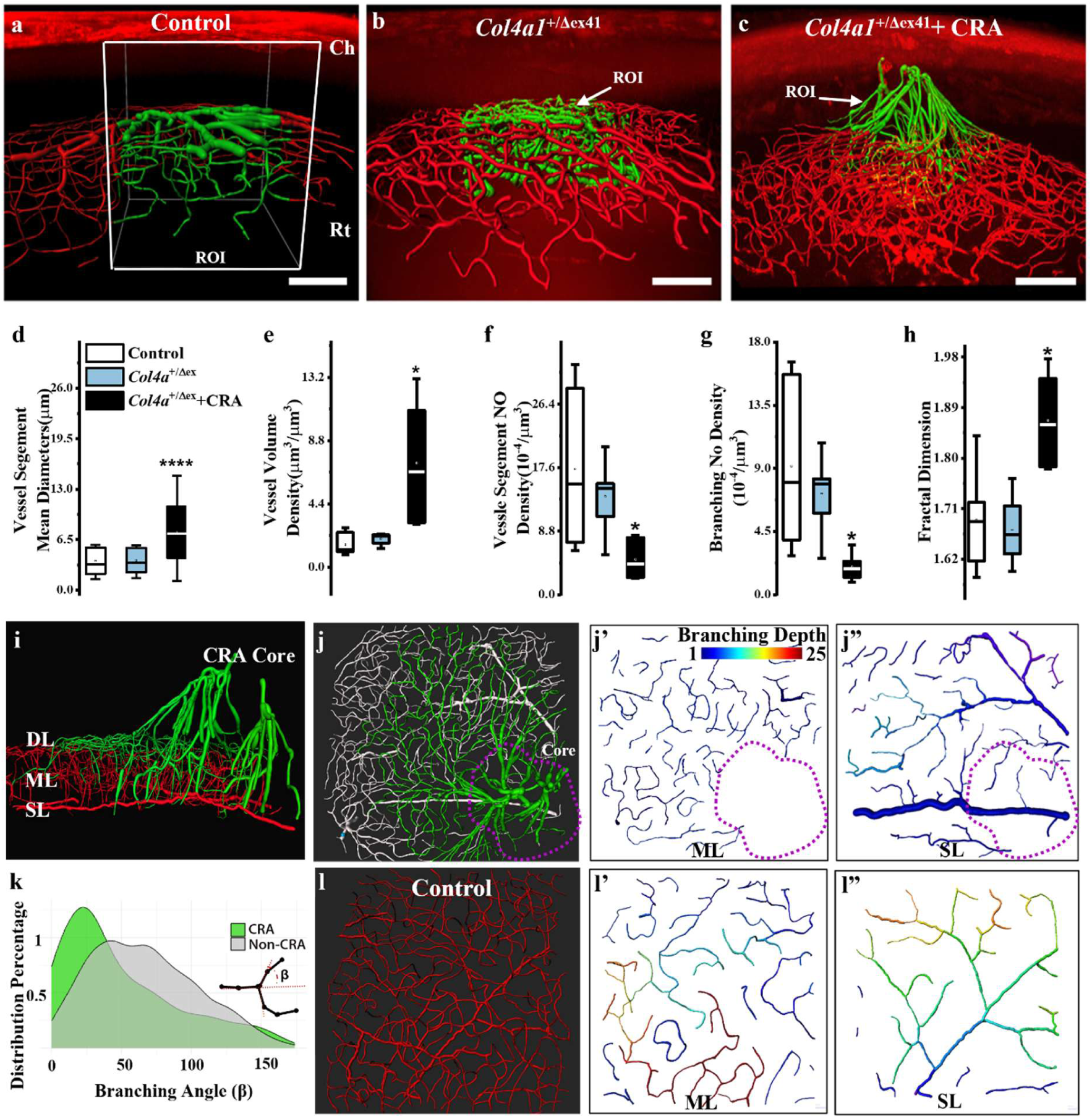
CRA lesions in Col4a1^+/Δex41^ mice are associated with capillary rarefaction, middle-layer loss, and radial vessel orientation. **(a)** Representative 3D reconstruction of control retina showing intact trilaminar vasculature organization. The white cube indicates the standardized 285 × 285 µm analysis window spanning the choroid (Ch) to the superficial layer plexus (SL). **(b)** Representative *Col4a1*^+/Δex41^ retina without neovascularization, demonstrating preserved laminar organization. **(c)** Representative *Col4a1*^+/Δex41^ retina with chorioretinal anastomosis (*Col4a1*^+/Δex41^ +CRA), showing disruption of trilaminar organization and aberrant vessels spanning the choroid and retina. For panels (a-c), vessels within ROI are highlighted in green. Scale bar, 100 µm. **(d)** Mean vessel segment diameter, **(e)** vessel-volume density, **(f)** vessel-number density, and **(g)** branching-point density, all quantified in full-thickness ROIs. **(h)** 3D fractal dimension (Bouligand–Minkowski method), reflecting network space-filling complexity. For analysis in panels **(d-h)**, one ROI was measured per retina per animal (n = 6 controls, n = 5 *Col4a1* without CRA, n = 6 *Col4a1*^+/Δex41^+CRA). Boxplots display the median (center line and interquartile range (Q1-3); whiskers represent 1.5 × IQR; outliers are shown individually. One-way ANOVA with Tukey’s post hoc test (* indicates *p* < 0.05; **** indicates *p* < 0.0001; ns, not significant). (i–j”) Characterization of CRA lesion. **(i)** Representative 3D reconstruction of a CRA illustrating layer-specific vascular alterations. The deep layer (DL) plexus extends abnormally toward and fuses with the choroid, the middle layer (ML) plexus shows regional depletion and discontinuity, and the superficial layer (SL) plexus appears relatively preserved. **(j)** En face projection of the same CRA lesion with demarcation of the CRA core by a purple dotted contour, wherein there is loss of the ML and radial convergence of vessels toward the lesion center. **(j’)** Branching-level map of the ML demonstrating significant network fragmentation, rarefaction, and absence of vessels within the CRA core. The color scale denotes number of connected branches within each grouping of vessels. **(j”)** Branching-level map of the SL within the same CRA demonstrating a continuous capillary network, with vessels oriented toward the lesion center. **(k)** Branching-angle analysis. Schematic defining bifurcation branching angle (β) defined as angle deviation from parent vessel trajectory. Within the CRA, the β distribution shifts toward smaller angles compared with non-CRA regions. (l-l’) Matched control retina. **(l)** En face projection of a littermate control retina showing a normal vascular network. **(l’)** Local branching-level map of the ML in a control retina, demonstrating normal network continuity and branching. **(l’’)** Local branching-level map of the SL in a control retina, demonstrating preserved network organization without radial orientation.

We identified two key structural domains in large CRA: a lesion core marked by loss of the retina’s trilaminar vascular architecture and aberrant choroid-to-retina vascular connections, and a surrounding peripheral region with disorganization of the retina’s three-layer network (Fig. 2i, j). Notably, the ML was completely absent within the CRA core and highly fragmented in the peripheral regions surrounding the lesion (Fig. 2j, k, l). Quantification also revealed that the ML fragmentation index in Col4a1^+/Δex41^ retinas (10.14 ± 1.63 fragments/mm²) was significantly elevated compared with controls (5.44 ± 1.20 fragments/mm²), representing an 86% increase (*p*<0.05; Fig. 3h). This pattern of ML disruption, with complete loss centrally and progressive fragmentation peripherally, suggests a spatial gradient of vulnerability radiating from the neovascular focus. CRA lesions also exhibited a distinctive tree-like configuration in which a small number of large choroidal anchor vessels gave rise to branches with radial orientation that integrated into all layers of the retinal vasculature (Fig. 2i). This architecture was accompanied by abnormally narrow vessel bifurcations, wherein branching angle (β) within CRA were significantly smaller than those in surrounding non-CRA vasculature (Fig. 2m). Notably, β of non-CRA vasculature in mutants did not differ significantly from controls, suggesting that the shift toward lower β values can provide a unique biomarker that helps identify and distinguish these aberrant neovessels as they merge into the pre-existing retinal plexuses.

**Figure 3.**
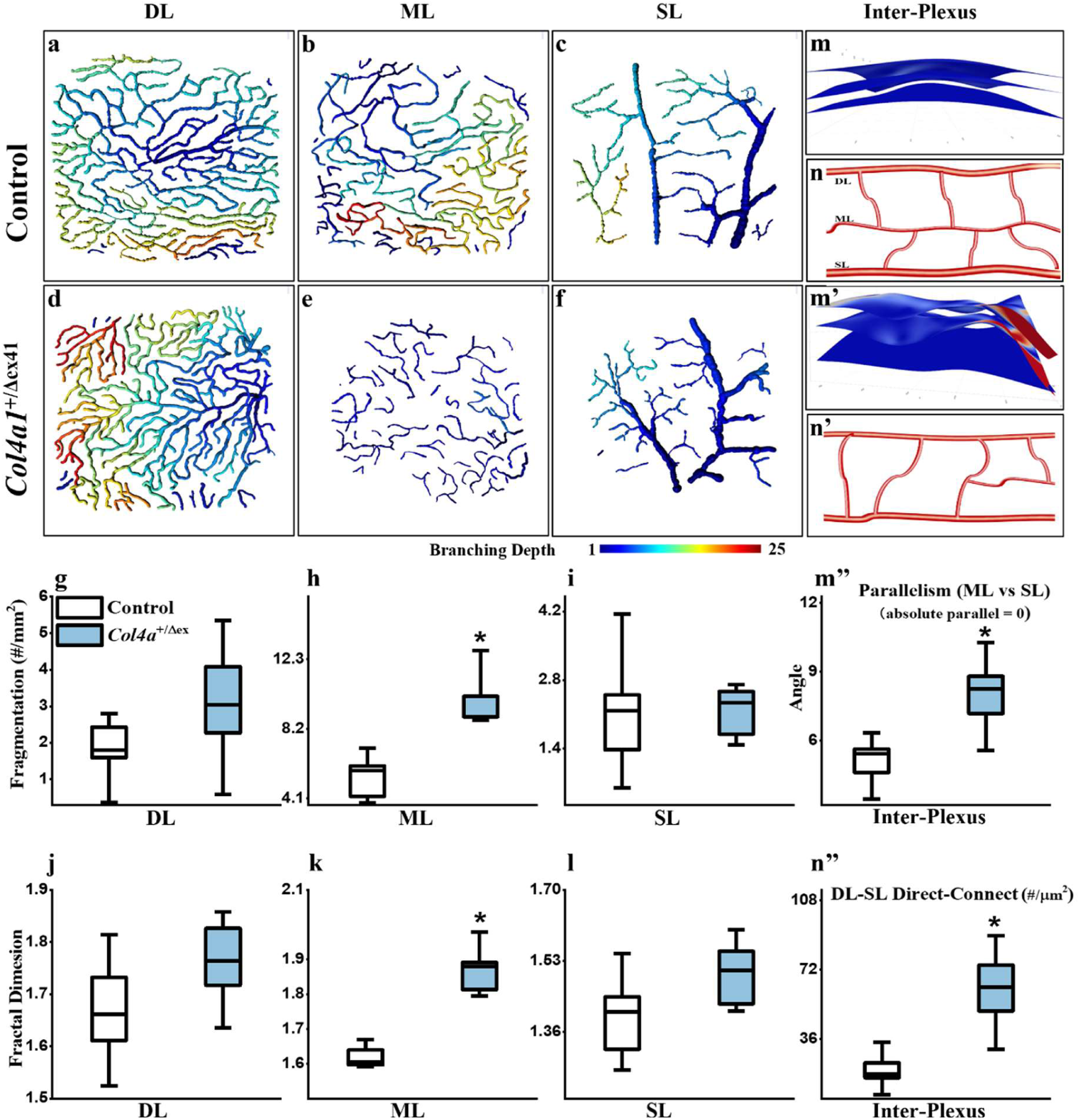
Middle-layer remodeling in Col4a1^+/Δex41^ mice without overt CRA. **(a–c)** Plexus-resolved vascular reconstructions from control retina showing deep (DL) (a), middle (ML) (b), and superficial (SL) (c) plexuses. All layers exhibit continuous, interconnected vessel architecture. Color scale denotes number of connecting branches within each grouping of vessels. **(d–f)** Corresponding plexus-resolved reconstructions from Col4a1^+/Δex41^ retina without CRA showing DL (d), ML (e), and SL (f). The ML exhibits discontinuity and fragmentation into isolated vascular segments, whereas DL and SL remain comparatively preserved. **(g–i)** Fragmentation index for DL (g), ML (h), and SL (i), defined as the number of connected vascular networks normalized to analyzed area (see Methods). ML fragmentation is significantly increased in Col4a1^+/Δex41^ retinas (h), whereas DL and SL show no significant change (g, i). **(j–l)** Plexus-specific 3D fractal dimension (FD) for DL (j), ML (k), and SL (l). ML FD is significantly altered in Col4a1^+/Δex41^ retinas (k), whereas DL and SL show no significant change (j,l). Box plots (g-l) show median and interquartile range (Q1–Q3); whiskers represent 1.5 × IQR; n = 6 control and n = 5 Col4a1^+/Δex41^ retinas; two-tailed Welch’s t-test; * indicates *p* < 0.05. **(m–m″)** Inter-plexus parallelism. **(m)** Representative point-wise parallelism heatmap from control retina showing alignment of ML and DL relative to SL, where cooler colors indicate greater parallelism (cosine similarity approaching 1). Spatial heatmaps and 3D surface visualizations are provided in Supplementary Fig. S2. **(m′)** Representative Col4a1^+/Δex41^ retina demonstrating reduced layer parallelism. **(m″)** Quantification of mean inter-plexus parallelism expressed as angular deviation from perfect alignment (0° = perfect parallel). ML–SL parallelism is significantly reduced in Col4a1^+/Δex41^ retinas; DL–SL parallelism is not significantly changed (n = 6 retinas per group; * indicates *p* < 0.05). **(n)** Schematic illustrating inter-plexus connections which typically include the ML in control retina. **(n′)** Schematic illustrating inter-plexus connections in Col4a1^+/Δex41^ retina with increased number of direct DL–SL connections. **(n″)** Number of DL–SL direct connections is substantially increased in Col4a1^+/Δex41^ retinas. Symbols indicate mean; error bars show ± standard error (SE) with caps indicating the 5th–95th percentiles; n = 6 control and n = 5 Col4a1^+/Δex41^ retinas; two-tailed Welch’s t-test; * indicates *p* < 0.05.

To identify architectural abnormalities that emerge before overt neovascular lesions form, we performed layer-resolved analysis comparing control and pre-neovascular Col4a1^+/Δex41^ retinas. Following manual stratification of the reconstructed networks into anatomically distinct layers (SL, ML, and DL) with comprehensive mapping of all inter-plexus connections (Fig. 3a–f, m’, n’), we conducted layer-specific 3D fractal dimension (FD) analysis and geometric quantification.

Initial 3D FD screening revealed that the ML and inter-plexus connections exhibited the most pronounced structural alterations. 3D FD of the ML was significantly elevated in Col4a1^+/Δex41^ retinas compared with controls (FD: 1.87 ± 0.07 vs. 1.61 ± 0.03, *p*<0.05; Fig. 3k). In contrast, the DL and SL showed no significant changes in fractal complexity (Fig. 3j, l), indicating that early structural remodeling is compartmentalized to the ML. Layer-level structural comparison revealed distinct connectivity alterations in the ML. While control ML networks formed continuous, hierarchically organized meshes with interconnected vessel segments (Fig. 3b), Col4a1^+/Δex41^ ML vessels appeared predominantly as disconnected fragments (Fig. 3e). This architectural disruption was quantitatively confirmed by 80% increase in ML fragmentation index (*p*<0.05; Fig. 3h), indicating substantial loss of network connectivity prior to neovascularization. Layer parallelism analysis revealed selective disruption of ML alignment in Col4a1^+/Δex41^ retinas. While control retinas maintained near-perfect parallel alignment across all layers (DL and ML deviation ∼5°; Fig. 3m), Col4a1^+/Δex41^ retinas showed significant ML misalignment (8.0 ± 1.9° vs 5.1 ± 1.2° in controls, *p*<0.05). Point-wise parallelism mapping revealed spatial heterogeneity in both layers (Fig. 3m, m’), but only the ML demonstrated statistically significant architectural disruption (Fig. 3m′′), identifying the ML as the primary site of early layer disorganization in pre-neovascular retinas. Quantification of inter-plexus connections revealed a disease-specific shift in cross-layer connectivity patterns. While DL-ML and SL-ML connections predominated in both groups, Col4a1^+/Δex41^ retinas showed a significant 3.5-fold increase in DL-SL direct connection density (63 ± 17 vs. 18 ± 10 connections per μm^2^; *p*<0.001; Fig. 3n″), with corresponding increases in vessel length per unit area. This compensatory re-routing between deep and superficial layers likely reflects adaptive responses to ML disruption. Together, these multi-scale analyses highlight the ML and inter-plexus connections as particularly sensitive compartments that undergo substantial architectural remodeling prior to overt neovascularization, with changes detectable through fragmentation, layer misalignment, and altered cross-layer connectivity patterns.

### Neuronal loss of Piezo2 disrupts 3D retinal vascular lattice development with selective alterations in the ML and inter-plexus connectivity

Neurovascular patterning of the retinal 3D vascular lattice depends on the coordinated development of planar vascular plexuses and the penetrating vessels that interconnect them. Our recent work with Toma et al^31^ established that a specialized subset of perivascular retinal ganglion cells (Nts-RGCs) guides the development of penetrating vessels through a Piezo2-dependent contact mechanism during retinal development. Motivated by this paradigm, we used Nts-RGC-specific Piezo2 knockout mice (Piezo2^Nts^) to evaluate whether our 3D quantification pipeline could sensitively detect developmental patterning defects and to determine how loss of neuron-derived mechanosensory cues alters plexus-resolved vascular architecture.

At the full-thickness scale, Piezo2^Nts^ retinas exhibited modest but significant increases in mean vessel diameter compared with controls (9.0 ± 2.0 vs. 6.8 ± 1.5 μm, *p*<0.05; Fig. 4b), with trends toward altered vessel volume density and branching point density (Fig. 4c, d). Full-thickness 3D fractal dimension (FD) analysis revealed a subtle but significant elevation (1.95 ± 0.03 vs. 1.91 ± 0.02, *p*<0.05; Fig. 4e), suggesting early increases in vascular surface irregularity and disruption of 3D lattice formation. Additional full-thickness metrics, including retinal thickness, convex hull volume, vessel segment count, total vessel length, mean segment length, and vessel straightness are provided in Supplementary Fig. S3.However, these full-thickness measurements obscured the compartment-specific architectural abnormalities that drive the phenotype, underscoring the critical importance of layer-resolved quantitative analysis.^35^ We separately evaluated SL, DL, and intermediate plexus (IP), where the IP encompasses the ML vessels and penetrating connections that develop and undergo active remodeling during the early postnatal period. Consistent with the temporal sequence of retinal vascular lattice assembly and the known role of Nts-RGCs in guiding penetrating vessel orientation, Piezo2 loss selectively and profoundly impaired IP organization while sparing the earlier-formed planar plexuses. Layer-resolved 3D FD analysis immediately identified the IP as the primary site of structural disruption. IP-specific FD was markedly elevated in Piezo2^Nts^ retinas (1.71 ± 0.04 vs. 1.63 ± 0.02, *p*<0.01; Fig. 4h), whereas DL and SL showed no significant changes (Fig. 4g, i). This compartmentalized complexity increase was accompanied by profound alterations in IP vessel geometry and connectivity. IP vessels displayed significantly larger mean segment diameters (8.5 ± 2.0 vs. 6.8 ± 1.5 μm, *p*<0.0001; Fig. 4h’) and a striking 40% increase in fragmentation index (1.28 ± 0.15 vs. 0.90 ± 0.15 ×10³ fragments/mm³, *p*<0.05; Fig. 4h”), indicating substantial disruption of intermediate layer network integrity. In contrast, while the SL and DL exhibited modest diameter increases, neither layer showed significant changes in network fragmentation or fractal complexity (Fig. 4g, i), supporting the interpretation that IP assembly represents a particularly vulnerable developmental program during the coordinated 2D to 3D vascular transition of the retina. Per-plexus analyses of additional morphometric parameters, including vessel segment count, branching points, total vessel length, mean segment length, and vessel straightness confirmed that these metrics did not differ significantly between genotypes in any layer (Supplementary Fig. S4), further supporting the specificity of IP disruption to fractal complexity, vessel diameter, and network fragmentation.

**Figure 4.**
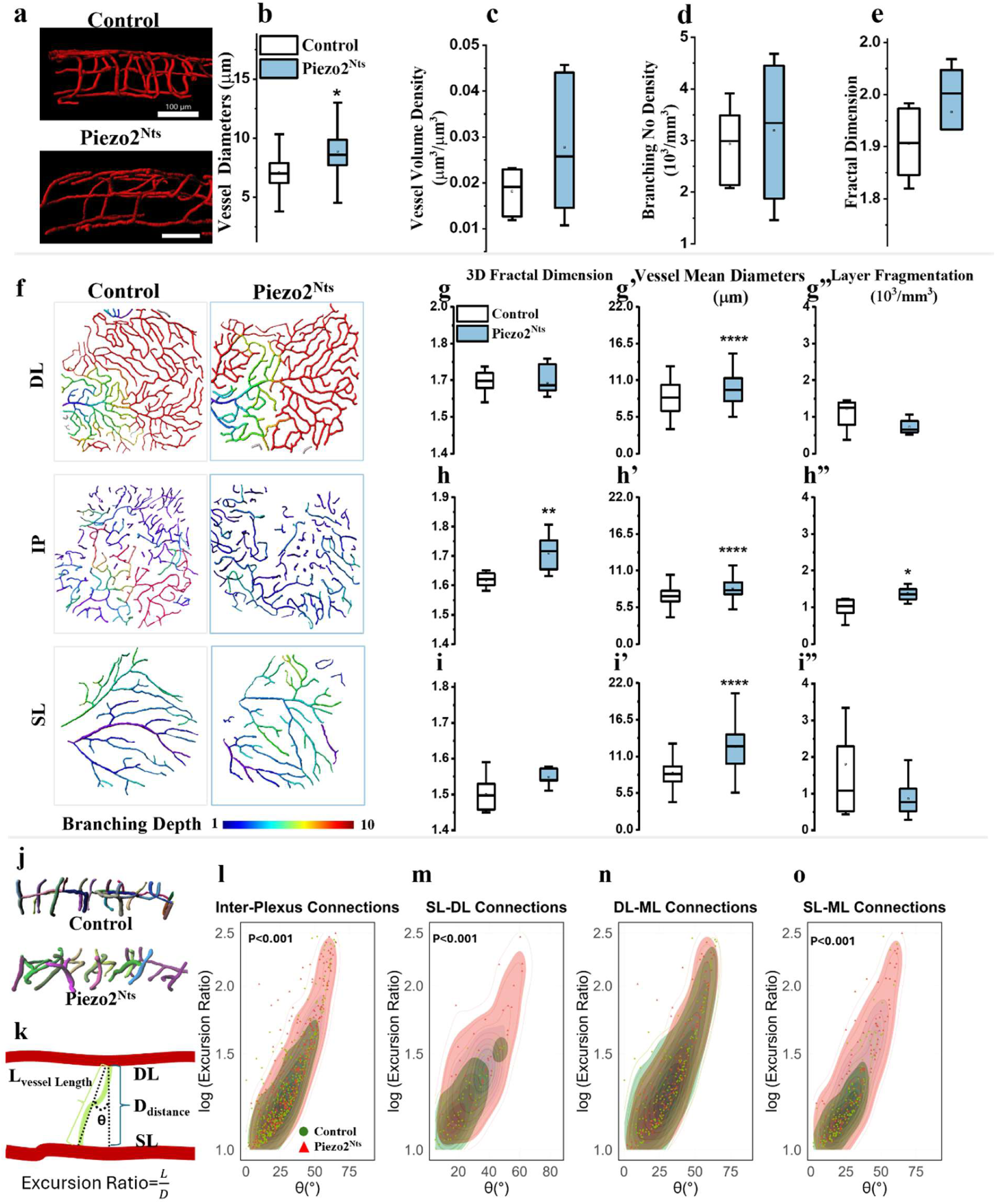
Neuron-specific loss of Piezo2 causes intermediate plexus fragmentation and altered penetrating vessel orientation and connectivity. **(a)** Representative side-view projections of full-thickness vasculature from Control and Piezo2^Nts^ retinas. While trilaminar vascular organization remains present in both groups, Piezo2^Nts^ retinas exhibit irregularities within the intermediate plexus. Scale bars, 100µm. **(b–e)** Full-thickness retinal analyses including: mean segment diameter (b), vessel-volume density (c), branching-point density (d), and 3D fractal dimension (FD; Bouligand–Minkowski method) (e). Full-thickness network analyses detect only modest but significant increases in mean diameter and global 3D FD of Piezo2^Nts^ retinas, with nonsignificant changes in branching and volume densities. Boxplots display median (center line), interquartile range (box), whiskers (1.5 × IQR), and individual outliers. n = 12 retinas per genotype (one Z-stack per retina). Two-tailed Welch’s t-test; *, *p* < 0.05; **, *p* < 0.01; ***, *p* < 0.001; ****, *p* < 0.0001. **(f)** Representative vascular reconstructions of the deep layer (DL) plexus, intermediate layer plexus (IP), and superficial layer (SL) plexus in Control and Piezo2^Nts^ retinas. Color scale denotes number of connecting branches within each grouping of vessels. **(g-g’’)** DL analyses including 3D FD (g), mean segment diameter (g’), and fragmentation index (g’’), demonstrating no statistically significant differences between Control and Piezo2^Nts^ retinas within the DL. **(h-h’’)** IP analyses including 3D FD (h), mean segment diameter (h’), and fragmentation index (h’’). The IP demonstrates higher 3D FD and mean segment diameter in Piezo2^Nts^ retinas compared with Controls. **(i-i’’)** SL analyses, including 3D FD (i), mean segment diameter (i’), and fragmentation index (i’’), showing no significant differences between groups. Boxplots as in (b–e). Two-tailed Welch’s t-test; *, *p* < 0.05; **, *p* < 0.01; ***, *p* < 0.001; ****, *p* < 0.0001. **(j)** Representative reconstructions of the IP highlighting altered penetrating vessel orientation in Piezo2^Nts^ retinas compared with Controls. **(k)** Schematic describing penetrating vessel characteristics including excursion ratio and orientation angle θ. **(l–o)** Joint distributions of log_10_(Excursion ratio) vs θ (degrees) for IP vessels including all IP vessels (l), SL–DL direct connections (m), DL–ML direct connections (n), and SL–ML direct connections (o). Piezo2^Nts^ retinas exhibit increased excursion ratio and larger θ, which are more pronounced among SL–ML and SL–DL and direct connecting vessels. *P*-values from Welch’s tests.

We further examined inter-layer connectivity by analyzing the geometry of penetrating vessels that join adjacent vascular plexuses. For each penetrating vessel, we defined the orientation angle (θ) as the deviation between the straight-line vector connecting its fusion points across plexuses and the axis orthogonal to the plexus planes (Fig. 4l, m). To quantify deviation from an ideal vertical trajectory, we calculated the excursion ratio (ER), defined as the ratio of the vessel’s true path length (L) to the orthogonal inter-plexus distance (D), where values approaching 1.0 indicate direct routing and higher values reflect tortuous or oblique trajectories. Piezo2^Nts^ retinas exhibited a pronounced shift toward oblique penetrating vessel trajectories. Mean orientation angles were significantly increased compared with controls (θ: 49.8 ± 10.7**°** vs. 27.6 ± 8.3°, *p* < 0.001), and penetrating vessels followed substantially longer and less direct paths, as reflected by elevated excursion ratios (log₁₀(ER): 1.91 ± 0.24 vs. 1.26 ± 0.17, *p* < 0.001; Fig. 4l). These changes indicate a transition from compact, near-orthogonal trajectories toward elongated and highly oblique courses spanning the inter-plexus space.

To evaluate how these geometric features covary, we paired θ and log₁₀(ER) for each penetrating vessel and constructed two-dimensional probability density distributions across inter-plexus classes (SL-DL, DL-ML, SL-ML, and combined). In Piezo2^Nts^ retinas, joint distributions were consistently shifted upward and rightward relative to controls, with density contours extending into higher-angle and higher-excursion regimes (θ ≈ 40–70°, log₁₀(ER) ≈ 1.6–2.3; Fig. 4l–o). In contrast, control distributions remained tightly concentrated within lower-angle, lower-excursion domains (θ ≈ 10–30°, log₁₀(ER) ≈ 1.0–1.4), consistent with well-aligned, vertically oriented penetrating vessels.

Stratification by connection subtype revealed that SL–DL and SL–ML vessels exhibited the most pronounced rightward and upward displacements in θ-ER space, together with broader contour spread, indicating both increased deviation and variability in penetrating vessel geometry (Fig. 4m, o). These connection types, originating from or terminating at the superficial plexus anatomically accessible to Nts-RGC somata and dendrites, displayed the most severe geometric perturbations and loss of stereotyped precision. In contrast, DL-ML connections showed comparatively smaller shifts (Fig. 4n), consistent with reduced Nts-RGC contact.

These findings demonstrate that Piezo2-dependent neuronal guidance is essential for establishing proper vertical scaffolding architecture and intermediate layer plexus organization during postnatal development. Notably, this IP-centered disruption in Piezo2^Nts^ retinas mirrors the vulnerability observed in the neovascularization model, where early ML disorganization precedes overt lesion formation. This convergence across developmental and pathological contexts underscores a shared susceptibility of the ML to distinct biological insults and highlights the sensitivity of our 3D quantitative framework for detecting common structural mechanisms underlying retinal vascular remodeling.

## Discussion

The neurovascular retina is a uniquely accessible extension of the central nervous system, enabling direct visualization of its vascular network and exquisitely organized laminar architecture. This combination of accessibility and structural precision has established the retina as a powerful model for studying vascular development and a promising window into systemic and neurovascular health. In this study, we establish a framework for detailed 3D retinal vascular quantification and demonstrate its ability to resolve overt pathology and subtle pre-pathological remodeling across distinct biological contexts. Applying this framework to models of spontaneous chorioretinal neovascularization in adult eyes and disrupted 3D neurovascular lattice formation during early retinal development, we provide quantitative insight into these processes and uncover a convergent vulnerability of the ML that emerges early, before overt neovascularization or architectural failures of the retinal vasculature.

Our approach demonstrates high sensitivity beyond conventional vessel-level metrics by resolving network and layer-specific architectural changes. In a pathological angiogenic setting, analysis of Col4a1^+/Δex41^ retinas enabled robust detection and characterization of spontaneous chorioretinal neovascularization, capturing hallmark features of disease, including vessel enlargement, capillary rarefaction, and increased morphologic complexity. Importantly, the framework also revealed architectural abnormalities extending beyond established lesion cores, including early fragmentation and misalignment of the ML and altered inter-plexus routing. These changes were also detectable prior to overt neovascularization, indicating that 3D microarchitectural disruption may represent an early marker of vascular instability rather than just a secondary consequence of formed lesions. The spatial gradient of ML loss, ranging from complete absence within lesion cores to progressive fragmentation in surrounding regions, supports a model in which failure of the middle plexus serves as an organizing feature of chorioretinal neovascular disease. This preferential vulnerability of the intermediate plexus aligns with emerging clinical evidence. Recent OCTA studies in diabetic retinopathy have demonstrated that the intermediate capillary plexus exhibits distinct and early density reductions that predict disease progression,^30^ and that capillary nonperfusion in the deeper plexuses is associated with both photoreceptor disruption and functional visual loss.^36^ These clinical observations, together with anatomical studies revealing the unique connectivity architecture of the IP in human retinas,^37^ support the notion that this vascular compartment occupies a structurally and functionally vulnerable position within the retinal vascular hierarchy. By contrast, during early retinal development, loss of Piezo2 in Nts-RGCs disrupted retinal vascular organization by impairing neurovascular guidance rather than causing pathological angiogenesis, producing a similarly profound but developmentally driven alteration of ML architecture. Although full-thickness metrics suggested only modest alterations, layer-resolved analysis revealed profound disruption of the intermediate plexus and its penetrating connections. Increased fragmentation, elevated fractal complexity, and geometric distortion of interplexus vessels point to failure of the vertical vascular scaffold that normally integrates planar plexuses into a coherent 3D lattice. These findings refine our prior work by demonstrating that Piezo2-dependent neuronal guidance is not merely required for penetrating vessel emergence, but is essential for preserving IP integrity during the critical developmental transition from planar to fully 3D vascular organization.

Oculomics is the quantitative assessment of ocular imaging features to derive imaging-based biomarkers of systemic, cardiovascular, and neurological disease.^38,39^ Prior studies of retinal vasculature have identified important indicators of systemic and neurological disorders using fundus photography and fluorescein angiography.^4,40^ More recently, OCT angiography has enabled depth-resolved visualization and separation of retinal vascular plexuses; however, most downstream quantitative analyses have relied on simplified 2D vascular representations and vessel metrics such as vessel caliber, density, tortuosity, and 2D fractal dimension.^41–43^ Deep learning approaches have further leveraged these representations, demonstrating encouraging performance for detecting biomarkers of cardiovascular^44^ and neurological disorders,^45,46^ including Alzheimer’s disease^47^ and cognitive impairment.^48,49^ Despite these advances, reliance on reduced 2D representations may obscure critical aspects of vascular network organization inherent to the retina’s 3D architecture. Leveraging fully 3D vascular representations, including layer-specific architecture and inter-plexus connectivity, therefore represents a critical opportunity to expand the biomarker space and advance the field of oculomics. In this study, we demonstrate that 3D, interlayer, and layer-specific analyses reveal subtle vascular alterations. We found the 3D fractal dimension to be an informative readout, as it implicitly captures network complexity, tortuosity, and layer curvature, which may be lost in 2D analyses.^50^ Our 3D analytical pipeline further enables the characterization of complex vascular phenotypes, such as CRAs, in which aberrant and neovascular structures traverse multiple retinal layers. This approach captures true spatial vascular trajectories and interlayer relationships, including the absence of the ML within CRA lesion cores and its fragmentation in the CRA periphery. Notably, both the Col4a1^+/Δex41^ and Piezo2^Nts^ mouse models evaluated in this study exhibit previously reported cerebrovascular abnormalities,^31,33,51,52^ supporting the link between retinal and cerebrovascular irregularities across disease contexts. Together, this work provides a framework for comprehensive 3D retinal vascular analysis that will facilitate future studies linkaging retinal microvascular organization with systemic and neurovascular pathology.

This study has several important limitations. Although the analytical framework we present is broadly applicable, the work described here presents a focused subset of analytical outputs, as we deliberately concentrated on metrics that were informative for the disease contexts examined. Several sub-analyses currently require manual parameter tuning, and vessel segmentation is only partially automated, necessitating manual verification and refinement; these steps can be time-intensive and highlight clear opportunities for increased automation, including deep learning-based segmentation and analysis pipelines to improve scalability and analysis throughput. Sample sizes were modest; however, these animal models and phenotypes were rigorously validated in our prior work,^31,33^ and statistically significant differences were consistently observed across groups, underscoring the sensitivity of the imaging and analytical framework. In addition, our analyses focused primarily on structural vascular features and did not incorporate physiological measurements such as blood flow or perfusion, which could further enhance biological interpretation and enable arterial-venous differentiation within the 3D framework. Finally, this study relied on advanced multiphoton imaging to achieve high-resolution, 3D visualization of the retinal vasculature, a modality not currently available in clinical practice. While this limits immediate translation to patients, it enabled the validation of high-fidelity analytical methods using subcellular-resolution datasets in preclinical models that can serve as structural ground truth for future adaptation to clinically accessible imaging modalities, particularly OCT angiography and emerging high-resolution 3D imaging technologies.^53^ As retinal imaging continues to evolve toward deeper volumetric analysis, the framework presented here provides a foundational platform that is readily adaptable to research and clinically deployable imaging systems.

This work establishes a comprehensive framework for 3D retinal vascular quantification that extends beyond conventional vessel-level metrics to reveal layer-specific vulnerabilities and network-level architectural principles governing vascular health and disease. Our discovery of convergent ML disruption across mechanistically distinct pathologies, including spontaneous chorioretinal neovascularization and impaired neurovascular patterning, suggests that middle-layer fragmentation may represent a universal structural signature of early retinal vascular instability across a wide range of inciting mechanisms. This interpretation is strengthened by clinical OCTA findings demonstrating that the intermediate capillary plexus shows early and differential vulnerability in diabetic retinopathy,^3,30^ and that deeper plexus nonperfusion predicts disease complications independently of superficial layer changes,^36^ pointing to a layer-specific hierarchy of vascular resilience that our 3D framework is uniquely positioned to resolve. By resolving true 3D spatial trajectories, inter-plexus connectivity patterns, and layer-specific complexity metrics, this framework captures microarchitectural features that are fundamentally inaccessible to 2D projection-based analyses yet may hold the key to earlier disease detection. The high-resolution anatomical maps generated here serve a dual purpose: first, as discovery tools for identifying which specific architectural features harbor the greatest diagnostic and prognostic value; and second, as quantitative benchmarks for developing algorithms that may detect these same features in lower-resolution but clinically available retinal angiography data. Immediate next steps include systematic validation of vascular biomarkers across expanded disease models, development of automated deep learning pipelines for 3D segmentation and analysis, and prospective evaluation of whether ML fragmentation and inter-plexus geometric distortion provide diagnostic or predictive information in human systemic and neurovascular diseases. As clinical retinal vascular imaging technologies continue their evolution toward higher-resolution volumetric acquisition, the methods and biological insights presented here provide a computational and conceptual foundation for incorporating layer-resolved retinal microvascular analysis into translational and clinical investigation.

## Methods

### Animals

All animal procedures were approved by the Laboratory Animal Resource Center at the University of California, San Francisco. Mice were housed in a pathogen-free facility under 12-h light/12-h dark cycles with ad libitum access to food and water. For the neovascular model, *Col4a1*^+/Δex41^ mice on C57BL/6J (B6) genetic background were crossed with 129 mice (009104; Jackson Laboratory), which develop spontaneous chorioretinal neovascularization and anastomoses in late adulthood, as previously described.^33^ Eyes of both sexes and littermate controls were harvested and prepared for imaging at 8-12 months of age. For the developmental model, Piezo2^Nts^ knockout C57BL/6J mice were used, which exhibit disrupted 3D retinal and cerebellar vascular lattice organization during early and perinatal development, as previously reported.^31^ Eyes were harvested at 2 months of age from both sexes as well as littermate controls.

### Retinal Vascular Imaging

Mice were anesthetized and transcardially perfused with phosphate-buffered saline (PBS). Whole eyes were harvested and fixed overnight in 4% paraformaldehyde (PFA) at 4°C. Eye specimens were depigmented using 3% H_2_O_2_ for 2 hours, then washed in PBS for 2 hours with gentle shaking at 37°C. Specimens were incubated with 5 μg/ml Lycopersicon esculentum (Tomato) lectin DyLight 649(Vector Laboratories, ZK0717) for 2 days at 37°C to label the vasculature. Labeled eye specimens were mounted in 1% agarose and imaged using a custom two-photon excited fluorescence microscope (modified Bergamo II, ThorLabs) using 1238 nm excitation light (Insight X3, SpectraPhysics) and a high numerical aperture water-dipping lens (Nikon 25× Apo LWD, 1.10 NA, 2.0 mm WD; THN25X-APO-MP1300) to acquire image data with subcellular resolution. To preserve native 3D retinal vascular architecture, image Z-stacks were acquired directly through the sclera of the intact globe using a 2-μm step size (Fig. 1a). Each Z-stack spanned approximately 358 μm in depth with a lateral pixel size of 1.12 μm, yielding a field of view of approximately 573 × 573 μm^2^. Interactive visualization of 3D vasculature and segmentation was performed using Imaris (10.2.0 Oxford Instruments).

### Vascular Network Segmentation

High-resolution image Z-stacks were stored in TIFF format with acquisition metadata. 3D vascular networks were reconstructed using a semi-automated segmentation pipeline comprising preprocessing, automated tracing, and manual validation steps. Raw images were preprocessed through sequential background subtraction, Gaussian filtering, baseline subtraction, median filtering, and deconvolution to enhance signal quality. Surface objects were generated to encompass vascular signal regions, creating masks that minimized noise artifacts. New imaging channels based on these masks were created and converted to the Imaris file format for analysis.

Automated vessel tracing was performed using the Imaris filament tracing algorithm. Traced structures were manually inspected to correct erroneous connections, add undetected vessels, and verify diameter measurements and branching connectivity. Validated 3D networks were manually stratified into anatomically distinct retinal layers: SL, ML, DL, and IP (Supplementary Video S1). Finalized networks were exported as VRML (wrl) 3D model files and accompanying CSV statistics files containing vessel segment lengths, diameters, branching coordinates, and positional data for downstream quantitative analysis. In the Imaris filament framework, vascular networks were represented as graph structures with vertices (discrete points along vessel centerlines) and edges (connections between vertices). Vertices were classified as terminal points (vessel endpoints), branching points (bifurcation or convergence nodes), or intermediate points (nodes along vessel segments). Segments were defined as collections of edges between branching or terminal vertices.

### Computational Analysis Pipeline

Following segmentation and validation in Imaris, downstream quantitative analysis was performed using Retina3D Vascular Analyzer, a hybrid Python/R framework designed for accessible and reproducible 3D vascular analysis. The pipeline requires minimal user setup: users export reconstructed 3D vascular networks from Imaris as VRML (.wrl) model files and CSV statistics files, rename them according to standardized naming conventions (with in-software guidance and documentation provided), and specify the input directory through the graphical interface to initiate analysis. The pipeline then operates in sequential stages: (1) geometry preprocessing, including convex hull computation (scipy.spatial.ConvexHull, Quickhull algorithm) and alpha shape analysis for volume normalization (Python: numpy, scipy, trimesh, pyvista); (2) three-dimensional fractal dimension analysis using the Bouligand–Minkowski morphological dilation method (Python); and (3) statistical analysis comprising full-thickness quantification, layer-resolved fractal analysis, and inter-layer connectivity assessment (R: mgcv, ggplot2, ggpubr, FSA, pracma). Each preprocessing module can also be used independently as a standalone tool for 3D geometric computation or fractal dimension calculation, enabling integration into other analytical workflows. The pipeline supports both interactive analysis through R Shiny dashboards and automated batch processing for reproducible HTML report generation. Complete source code, installation instructions, example datasets, and interactive documentation are available at the project repository (see Code Availability).

### Vascular Network Quantification and Analysis

Vascular network architecture was characterized using multiple morphometric and topological parameters derived from validated 3D models (Table 1). All quantitative analyses were performed within standardized 285 × 285 μm regions of interest (ROI), spanning full retinal thickness from choroid to the SL. The ROI dimension represents the x- and y-axis half-widths of the raw data, and regions were selected to avoid edge artifacts and minimize peripheral optical aberrations of the acquisition window.

#### Vessel density parameters

Vessel number density and branching point density were calculated by normalizing total counts to the ROI or full imaging region convex hull volume. Vessel volume density was calculated by dividing the total network volume by the ROI or full imaging region convex hull volume. Convex hull volumes were computed from the 3D vascular point cloud using the Quickhull algorithm (scipy.spatial.ConvexHull), providing a standardized tissue occupancy volume for density normalization. These density metrics provide complementary information: number and branching point densities reflect vascular branching complexity, while volume density captures the total vascular space occupancy within the tissue.

#### Branching architecture and hierarchical organization

Branch levels were assigned using diameter-based classification. Starting from vessel origins (Branch Level 1), segments at each bifurcation were classified hierarchically: smaller-diameter daughter segments received incremented branch levels, while larger-diameter segments maintained parent levels, capturing network organization from major vessels to terminal capillaries. Branching angles were quantified as angular deviation between vessel segments and parent vessel extension direction (Supplementary Fig. S5), reflecting local bifurcation geometry. Specifically, α represents the angle between the two daughter vessel segments extending from the bifurcation point to their distal endpoints, while β captures the angular deviation between the parent vessel’s extended centerline direction and each daughter vessel’s initial segment (β_1_ and β_2_), measured at the nearest vertices from the bifurcation point. Together, these metrics characterize both the hierarchical structure and geometric properties of vascular branching patterns, enabling comparative analysis across developmental and neovascular contexts.

#### Inter-plexus parallelism

Spatial organization between vascular layers was quantified using Generalized Additive Models (GAM) with 3D surface analysis. 3D point clouds representing vessel positions were extracted from each vascular layer (minimum 8 points per layer due to the algorithm requirement). For each layer, a GAM with thin-plate regression splines fitted smooth surfaces to the positional data. Surface normal vectors were computed at 150×150 grid point using numerical gradient estimation followed by L2 normalization. Parallelism between layers was quantified using cosine similarity between corresponding normal vectors, with the superficial layer serving as reference. GAM fitting was performed using the mgcv package in R.^54^ Group differences were assessed using non-parametric tests (Wilcoxon rank-sum for two groups, Kruskal-Wallis for multiple groups) appropriate for angular measurements.

#### Inter-plexus vessel geometry

For vessels connecting retinal layers, two complementary geometric parameters were quantified. Vessel orientation angles (θ) were calculated relative to the superficial layer reference frame using local vector analysis at each segment (Fig. 4k), characterizing 3D routing patterns and cross-layer connectivity. Excursion ratio was defined as vessel path length divided by vertical span between layers, with values approaching 1 indicating direct perpendicular connections and higher values reflecting more tortuous routing. Detailed computational procedures are provided in Supplementary Methods S.M.1.

#### Fractal dimension analysis

3D fractal dimension (3D FD) was computed using the Bouligand–Minkowski morphological dilation method. Vascular objects were iteratively dilated using spheres of increasing radius *r* (voxel size = 1 unit, corresponding to 0.2 μm absolute resolution based on analyzed mesh resolution), and the resulting influence volume V(r) was computed at each dilation step. The fractal dimension was derived from the negative slope of the linear regression of log V(r) versus log r within the scale-invariant interval (Equation 1, Supplementary Fig. S6). FD was computed using two complementary geometric representations: (1) skeletonized centerlines, capturing branching topology independent of vessel diameter, and (2) surface meshes, reflecting volumetric occupancy and surface irregularity. This dual approach enabled distinction between topological complexity and spatial occupancy. By definition, FD ranges from 0 to 3, with higher values indicating greater structural complexity; values for retinal vascular networks in this study ranged from approximately 1.6 to 2.0. The Bouligand–Minkowski method was selected for its robustness and sensitivity to subtle architectural changes in biological networks, demonstrating close agreement with theoretical FD values in mathematical benchmarks.^55^

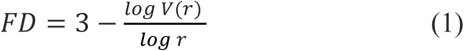

### Statistical Methods

All quantitative analyses were performed on 3D geometric data (CSV files) and reconstructed retinal 3D model files (WRL files) exported from Imaris. Data processing and statistical analyses were implemented in Python 3.10 (with numpy, scipy, trimesh, and pyvista for geometric and fractal computations) and R 4.4.0 (with tidyverse, ggplot2, ggpubr, mgcv, FSA, and pracma for statistical analysis and visualization) using custom scripts (see code availability).

Quantitative readouts included vessel segment length, diameter, tortuosity, branching order, branching angles, vessel density, and fractal dimension. For fractal dimension analysis, both skeletonized centerlines and vascular surface meshes were evaluated using the Bouligand–Minkowski dilation method, enabling complementary assessment of topological versus volumetric complexity. Inter-plexus geometry was assessed by quantifying parallelism and direct bridging connections between plexuses. Complete definitions of all quantification parameters available for retinal vessel network evaluation are provided in Table 1.

Statistical comparisons were conducted using R base and the stats package, for parametric comparisons between two groups, two-tailed Welch’s t-tests were applied after verifying normality using Shapiro-Wilk tests. Multi-group comparisons were performed using one-way ANOVA with Tukey’s honest significant difference post hoc tests to control for multiple comparisons. Nonparametric alternatives (Wilcoxon rank-sum test for two groups, Kruskal-Wallis test with Dunn’s post hoc for multiple groups) were used when normality assumptions were violated or when distributions were non-Gaussian for angular measurements.

Sample sizes for each experimental group are reported in the figure legends. All statistical tests were two-sided, with a significance threshold of α = 0.05. No adjustments were made for multiple comparisons across different vascular parameters, as each represents an independent biological measurement. Data are presented as mean ± standard deviation unless otherwise noted. Boxplots display median (center line), interquartile range (box bounds), whiskers extending to 1.5× IQR, and outliers as individual points beyond whiskers.

Sample sizes were determined based on effect sizes observed in previous studies of retinal vascular pathology and developmental models using these genetic lines,^31,33^ and were sufficient to detect statistically significant differences across all primary endpoints. All experiments included littermate controls to minimize genetic background effects.

### Vessel Network Terminology

Retinal vascular networks were represented as graph-based filament structures following IMARIS conventions, where vessels are decomposed into vertices (discrete points along vessel centerlines) and edges (connections between adjacent vertices). Vertices were classified by topological position as terminal points (vessel endpoints), branching points (bifurcation or convergence nodes), or intermediate points (nodes along vessel segments), with segments defined as collections of edges between branching or terminal vertices representing individual vessel branches within the network hierarchy. For anatomically specific analyses, vessels were stratified into distinct retinal compartments: superficial layer plexus (SL, ganglion cell layer), middle layer plexus (ML, intermediate capillary plexuses), deep layer plexus (DL, outer plexiform layer), and intermediate plexus (IP, ML with perpendicular vessels linking retinal layers). This graph-based representation enabled quantitative morphological and topological analysis of complex 3D vascular architectures.

## Data Availability

Example datasets for pipeline validation are included in the Retina3D Vascular Analyzer pipeline repository. Additional multiphoton imaging data and processed vascular network models from this study are available upon request.

## Code Availability

The Retina3D Vascular Analyzer pipeline (v1.1) is freely available at https://ucsf.box.com/v/Retina3DVascularAnalyzer. The repository includes complete installation guides, example datasets, and interactive user manuals.

## Author Contributions

W.S. designed the study, collected and analyzed the data, developed the computational analysis framework, and wrote the manuscript. G.H. assisted with experiments and data collection. W.E.K. assisted with data analysis and edited the manuscript. R.A.M. helped with analysis and editing the manuscript. P.Z. helped with analysis and editing the manuscript. K.T. collected data and edited the manuscript. X.D. provided key materials, result interpretation, and edited the manuscript. D.B.G. provided key materials, result interpretation, and edited the manuscript. T.N.K. conceived and supervised the study, interpreted results, and wrote the manuscript.

## Acknowledgements

We thank Dr. Frank Tahvi, Mr. Arjun Shivkumar, Dr. Cassandre Labelle-Dumais and Dr. Mao Mao for their intellectual input and discussions. This work was made possible by National Institutes of Health (NEI-K08EY033030, NEI-K12EY031372, NEI-P30EY002162, R01EY030138), BrightFocus Foundation (M2021015N), Research to Prevent Blindness, as well as generous support from the All May See Foundation, Huang Pacific Foundation, and Dr. David and Cecilia Lee.

## Competing Interests

The authors declare no competing interests.

## Notes

### Competing Interest Statement

The authors have declared no competing interest.

### Summary of Updates

Figure 1 Updated terminology from"Whole-strucure" to " Full-thickness".

## References

1. Fruttiger, M. Development of the retinal vasculature. Angiogenesis 10, 77–88 (2007).

2. Patton, N. et al. Retinal vascular image analysis as a potential screening tool for cerebrovascular disease: a rationale based on homology between cerebral and retinal microvasculatures. J. Anat. 206, 319–348 (2005).

3. Fayed, A. E. et al. Retinal vasculature of different diameters and plexuses exhibit distinct vulnerability in varying severity of diabetic retinopathy. Eye 38, 1762–1769 (2024).

4. Poplin, R. et al. Prediction of cardiovascular risk factors from retinal fundus photographs via deep learning. *Nat*. Biomed. Eng. 2, 158–164 (2018).

5. Geerling, C. F. et al. Changes of the retinal and choroidal vasculature in cerebral small vessel disease. Sci. Rep. 12, 3660 (2022).

6. Cheung, C. Y. et al. A deep-learning system for the assessment of cardiovascular disease risk via the measurement of retinal-vessel calibre. *Nat*. Biomed. Eng. 5, 498–508 (2021).

7. Danielescu, C. et al. Automated Retinal Vessel Analysis Based on Fundus Photographs as a Predictor for Non-Ophthalmic Diseases—Evolution and Perspectives. J. Pers. Med. 14, 45 (2023).

8. Liu, Y. et al. AI-based 3D analysis of retinal vasculature associated with retinal diseases using OCT angiography. Biomed. Opt. Express 15, 6416–6432 (2024).

9. Spaide, R. F., Fujimoto, J. G., Waheed, N. K., Sadda, S. R. & Staurenghi, G. Optical coherence tomography angiography. Prog. Retin. Eye Res. 64, 1–55 (2018).

10. Sampson, D. M., Dubis, A. M., Chen, F. K., Zawadzki, R. J. & Sampson, D. D. Towards standardizing retinal optical coherence tomography angiography: a review. Light Sci. Appl. 11, 63 (2022).

11. Zahid, S. et al. Fractal Dimensional Analysis of Optical Coherence Tomography Angiography in Eyes With Diabetic Retinopathy. Invest. Ophthalmol. Vis. Sci. 57, 4940–4947 (2016).

12. Durbin, M. K. et al. Quantification of Retinal Microvascular Density in Optical Coherence Tomographic Angiography Images in Diabetic Retinopathy. JAMA Ophthalmol. 135, 370–376 (2017).

13. Zhang, M. et al. Projection-resolved optical coherence tomographic angiography. Biomed. Opt. Express 7, 816–828 (2016).

14. Kim, T. N. et al. Line-Scanning Particle Image Velocimetry: An Optical Approach for Quantifying a Wide Range of Blood Flow Speeds in Live Animals. PLoS ONE 7, e38590 (2012).

15. Wang, T. Three-photon imaging of mouse brain structure and function through the intact skull. Nat. Methods 15, 9 (2018).

16. Qin, Z. et al. Adaptive optics two-photon microscopy enables near-diffraction-limited and functional retinal imaging in vivo. Light Sci. Appl. 9, 79 (2020).

17. Shih, A. Y. et al. Two-Photon Microscopy as a Tool to Study Blood Flow and Neurovascular Coupling in the Rodent Brain. J. Cereb. Blood Flow Metab. 32, 1277–1309 (2012).

18. Zhang, J. et al. Optical sectioning methods in three-dimensional bioimaging. Light Sci. Appl. 14, 11 (2025).

19. Rubart, M. Two-photon microscopy of cells and tissue. Circ. Res. 95, 1154–1166 (2004).

20. Zhang, Q. et al. Retinal microvascular and neuronal pathologies probed in vivo by adaptive optical two-photon fluorescence microscopy. eLife 12, e84853 (2023).

21. Hirano, T., Chanwimol, K., Weichsel, J., Tepelus, T. & Sadda, S. Distinct Retinal Capillary Plexuses in Normal Eyes as Observed in Optical Coherence Tomography Angiography Axial Profile Analysis. Sci. Rep. 8, 9380 (2018).

22. Tahir, W. et al. Anatomical Modeling of Brain Vasculature in Two-Photon Microscopy by Generalizable Deep Learning. BME Front. 2020, 8620932 (2020).

23. Zhang, J. et al. 3D microvascular reconstruction in retinal OCT angiography images via domain-adaptive learning. Pattern Recognit. 165, 111494 (2025).

24. Garrity, S. T., Iafe, N. A., Phasukkijwatana, N., Chen, X. & Sarraf, D. Quantitative Analysis of Three Distinct Retinal Capillary Plexuses in Healthy Eyes Using Optical Coherence Tomography Angiography. Investig. Opthalmology Vis. Sci. 58, 5548 (2017).

25. Zudaire, E., Gambardella, L., Kurcz, C. & Vermeren, S. A Computational Tool for Quantitative Analysis of Vascular Networks. PLOS ONE 6, e27385 (2011).

26. Narotamo, H., Silveira, M. & Franco, C. A. 3DVascNet: An Automated Software for Segmentation and Quantification of Mouse Vascular Networks in 3D. Arterioscler. Thromb. Vasc. Biol. 44, 1584–1600 (2024).

27. Bumgarner, J. R. & Nelson, R. J. Open-source analysis and visualization of segmented vasculature datasets with VesselVio. *Cell Rep*. Methods 2, 100189 (2022).

28. Milde, F., Lauw, S., Koumoutsakos, P. & Iruela-Arispez, M. L. The mouse retina in 3D: quantification of vascular growth and remodeling. Integr. Biol. Quant. Biosci. Nano Macro 5, 1426–1438 (2013).

29. Stahl, A. et al. The Mouse Retina as an Angiogenesis Model. Invest. Ophthalmol. Vis. Sci. 51, 2813–2826 (2010).

30. Haddad, C., Baleine, M. & Motulsky, E. An OCT-A Analysis of the Importance of Intermediate Capillary Plexus in Diabetic Retinopathy: A Brief Review. J. Clin. Med. 13, 2516 (2024).

31. Toma, K. et al. Perivascular neurons instruct 3D vascular lattice formation via neurovascular contact. Cell 187, 2767–2784.e23 (2024).

32. Selvam, S., Kumar, T. & Fruttiger, M. Retinal vasculature development in health and disease. Prog. Retin. Eye Res. 63, 1–19 (2018).

33. Alavi, M. V. et al. Col4a1 mutations cause progressive retinal neovascular defects and retinopathy. Sci. Rep. 6, 18602 (2016).

34. Lian, Y., Li, G., Liu, H. & Zhang, Q. Fractal analysis of retinal microvasculature and cardiovascular outcome: a systematic review and meta-analysis. BMC Ophthalmol. 25, 668 (2025).

35. Chang, C.-C. et al. Three-dimensional Imaging Coupled with Topological Quantification Uncovers Retinal Vascular Plexuses Undergoing Obliteration. Theranostics 11, 1162–1175 (2021).

36. Ong, J. X., Konopek, N., Fukuyama, H. & Fawzi, A. A. Deep Capillary Nonperfusion on OCT Angiography Predicts Complications in Eyes with Referable Nonproliferative Diabetic Retinopathy. *Ophthalmol*. Retina 7, 14–23 (2023).

37. Nesper, P. L. & Fawzi, A. A. Human Parafoveal Capillary Vascular Anatomy and Connectivity Revealed by Optical Coherence Tomography Angiography. Invest. Ophthalmol. Vis. Sci. 59, 3858–3867 (2018).

38. Zhu, Z. et al. Oculomics: Current concepts and evidence. Prog. Retin. Eye Res. 106, 101350 (2025).

39. Wagner, S. K. et al. Insights into Systemic Disease through Retinal Imaging-Based Oculomics. Transl. Vis. Sci. Technol. 9, 6 (2020).

40. Wong, T. Y. et al. Retinal microvascular abnormalities and 10-year cardiovascular mortality: A population-based case-control study. Ophthalmology 110, 933–940 (2003).

41. Liu, S. et al. Beyond the eye: A relational model for early dementia detection using retinal OCTA images. Med. Image Anal. 102, 103513 (2025).

42. Gao, Y. et al. A narrative review of retinal vascular parameters and the applications (Part I): Measuring methods. Brain Circ. 9, 121–128 (2023).

43. Campbell, J. P. et al. Detailed Vascular Anatomy of the Human Retina by Projection-Resolved Optical Coherence Tomography Angiography. Sci. Rep. 7, 42201 (2017).

44. Arnould, L. et al. Using Artificial Intelligence to Analyse the Retinal Vascular Network: The Future of Cardiovascular Risk Assessment Based on Oculomics? A Narrative Review. Ophthalmol. Ther. 12, 657–674 (2023).

45. Zhang, J., Shi, L. & Shen, Y. The retina: A window in which to view the pathogenesis of Alzheimer’s disease. Ageing Res. Rev. 77, 101590 (2022).

46. Yao, J., Hong, A. S. Y., Fukutsu, K. & Ting, D. S. W. Artificial intelligence oculomics for systemic health and longevity medicine: 2025 and beyond. Curr. Opin. Ophthalmol. 36, 477–486 (2025).

47. Cheung, C. Y. et al. Retinal imaging in Alzheimer’s disease. J. Neurol. Neurosurg. Psychiatry 92, 983–994 (2021).

48. Haft-Javaherian, M. et al. Deep convolutional neural networks for segmenting 3D in vivo multiphoton images of vasculature in Alzheimer disease mouse models. PloS One 14, e0213539 (2019).

49. Xie, J. et al. Deep segmentation of OCTA for evaluation and association of changes of retinal microvasculature with Alzheimer’s disease and mild cognitive impairment. Br. J. Ophthalmol. 108, 432–439 (2024).

50. Krohn, S. et al. Evaluation of the 3D fractal dimension as a marker of structural brain complexity in multiple-acquisition MRI. Hum. Brain Mapp. 40, 3299–3320 (2019).

51. Labelle-Dumais, C. et al. COL4A1 Mutations Cause Ocular Dysgenesis, Neuronal Localization Defects, and Myopathy in Mice and Walker-Warburg Syndrome in Humans. PLOS Genet. 7, e1002062 (2011).

52. Kuo, D. S., Labelle-Dumais, C. & Gould, D. B. COL4A1 and COL4A2 mutations and disease: insights into pathogenic mechanisms and potential therapeutic targets. Hum. Mol. Genet. 21, R97–R110 (2012).

53. Wang, L., Wang, B., Chhablani, J., Sahel, J. A. & Pi, S. Freqformer: Frequency-Domain Transformer for 3-D Reconstruction and Quantification of Human Retinal Vasculature. IEEE Trans. Biomed. Eng. 1–11 (2025) doi:10.1109/TBME.2025.3612332.

54. Wood, S. N. Fast Stable Restricted Maximum Likelihood and Marginal Likelihood Estimation of Semiparametric Generalized Linear Models. J. R. Stat. Soc. Ser. B Stat. Methodol. 73, 3–36 (2011).

55. Backes, A. R. & Bruno, O. M. Fractal and Multi-Scale Fractal Dimension analysis: a comparative study of Bouligand-Minkowski method. Preprint at 10.48550/arXiv.1201.3153 (2012).

