## Supplementary for "A Three-dimensional Analytical Framework for Retinal Microvasculature Reveals Layer-associated Vulnerability in Development and Neovascular Remodeling"

for

Wenhao Shang et al.

**This file contains:**

Supplementary Figures S1–S6

Supplementary Video S1-3

Supplementary Methods S.M.1–S.M.3

### Supplementary Figures

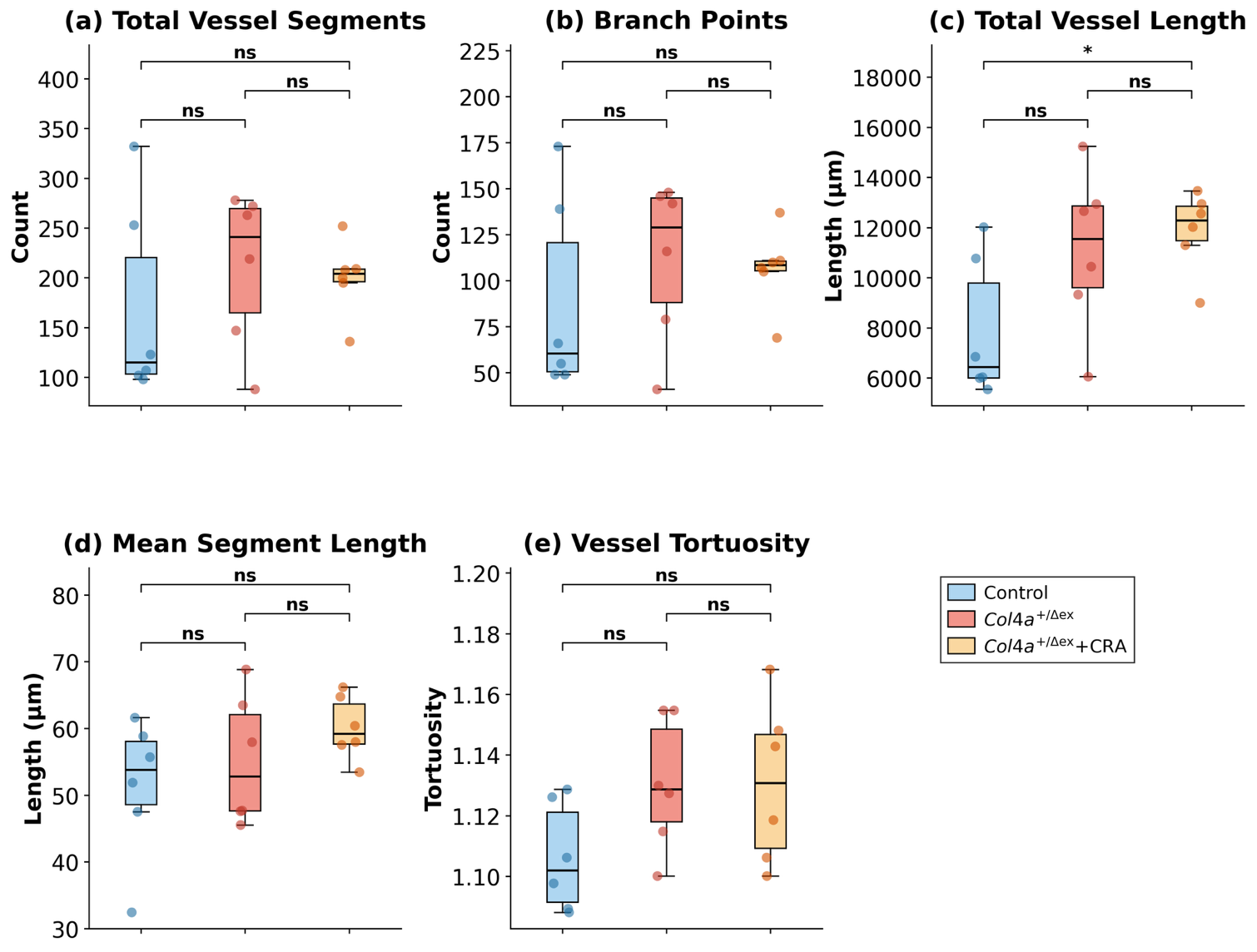

**Supplementary Figure S1. Additional vascular morphometric analyses in Col4a1**^+/Δex41^ **retinas with and without CRA lesions.** (a) Total number of vessel segments per region of interest (ROI). (b) Total branching point count per ROI. (c) Total vessel length (sum of all segment lengths) per ROI. CRA-containing retinas showed significantly greater total vessel length compared to controls (*p* < 0.05) (d) Mean segment length per ROI. (e) Vessel tortuosity (vessel length / end-to-end distance). No significant differences in segment count, branching, mean segment length, or tortuosity were observed between groups. Boxplots display the median (center line) and interquartile range (IQR, box boundaries); whiskers extend to 1.5× IQR. Statistical comparisons were performed using two-tailed Welch’s t-tests; **p* < 0.05, ns = not significant.

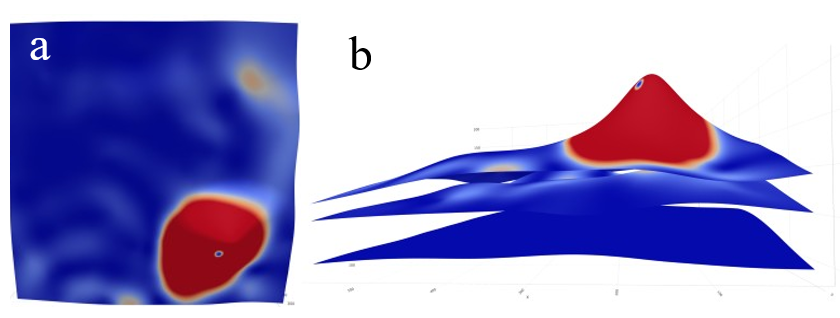

**Supplementary Figure S2. Inter-plexus parallelism analysis of retinal vascular layers in chorioretinal anastomosis.** (a) Heatmap showing the cosine value of the angular deviation between generalized additive model (GAM)-fitted surfaces of paired vascular layers（DL vs SL）. The color scale represents the local angle cosine (degrees) between surface normal vectors; warmer colors indicate greater deviation (lower parallelism). (b) Three-dimensional surface visualization of the GAM-fitted plexus surfaces for both layers.

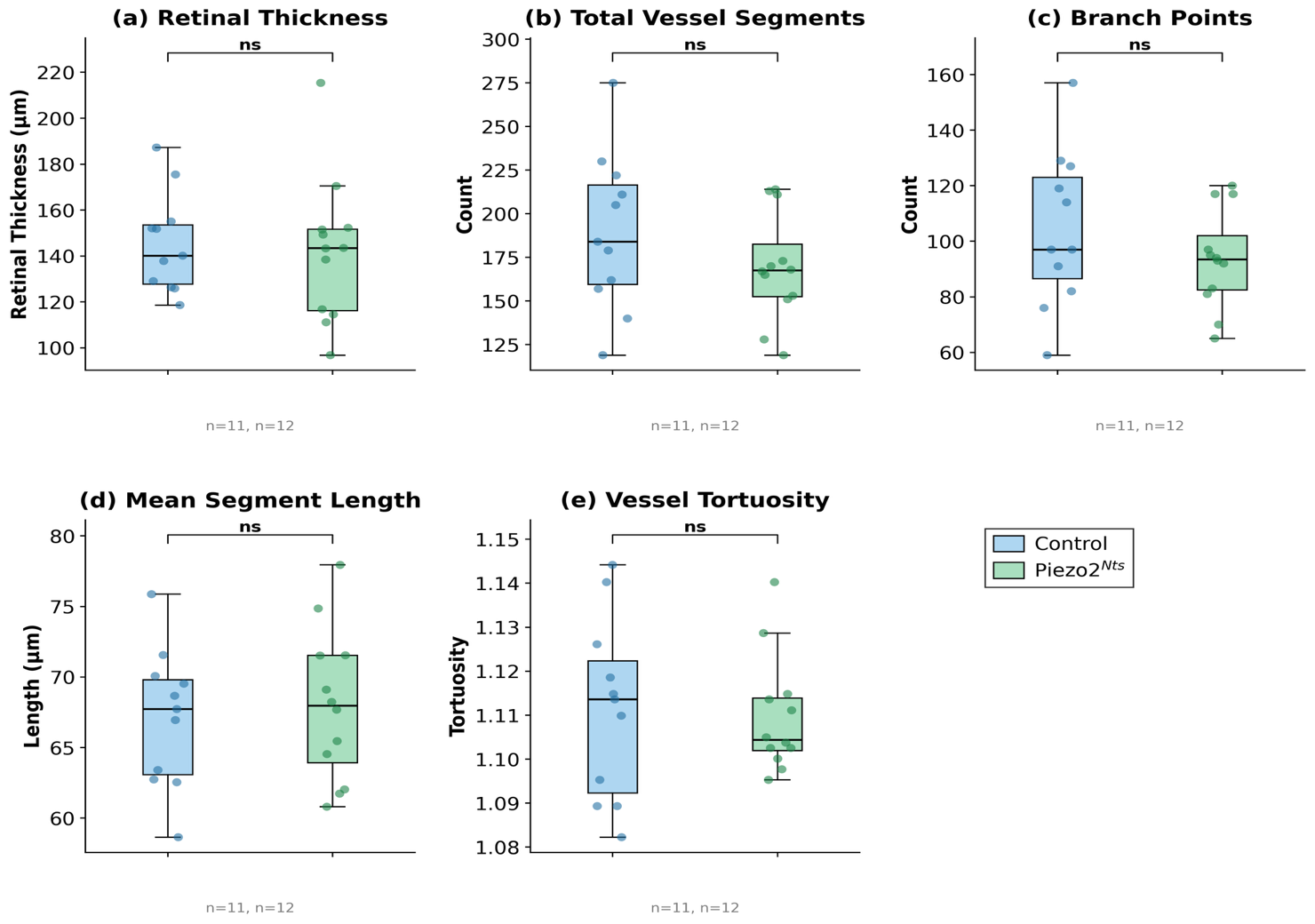

**Supplementary Figure S3. Additional vascular morphometric analyses of Piezo2^Nts^ retinas.**

(a) Retinal thickness, measured as the distance of the DL and SL. Control and Piezo2^Nts^ retinas showed comparable thickness (Control: 145.4 ± 21.6 μm; Piezo2Nts: 142.0 ± 31.4 μm; *p* = 0.76) (b) Total vessel segment count and (c) total branching point count. Both metrics showed a modest, non-significant reduction in Piezo2^Nts^ retinas (segments: −10.6%; branch points: −10.3%), suggesting a trend toward reduced vascular complexity that did not reach statistical significance at the whole-structure level. (d) Mean segment length, representing the average inter-branch distance within the vascular network; no difference was observed between groups (*p* = 0.68). (e) Vessel tortuosity (vessel length divided by end-to-end distance; values > 1 indicate increasing curvature); both groups exhibited similar tortuosity profiles (*p* = 0.83). Control (Ctrl, n = 11) and Piezo2^Nts^ (Nts, n = 12). Boxplots display the median (center line) and interquartile range (IQR, box boundaries); whiskers extend to 1.5× IQR. Statistical comparisons: two-tailed Welch’s t-tests; **p* < 0.05, ns = not significant.

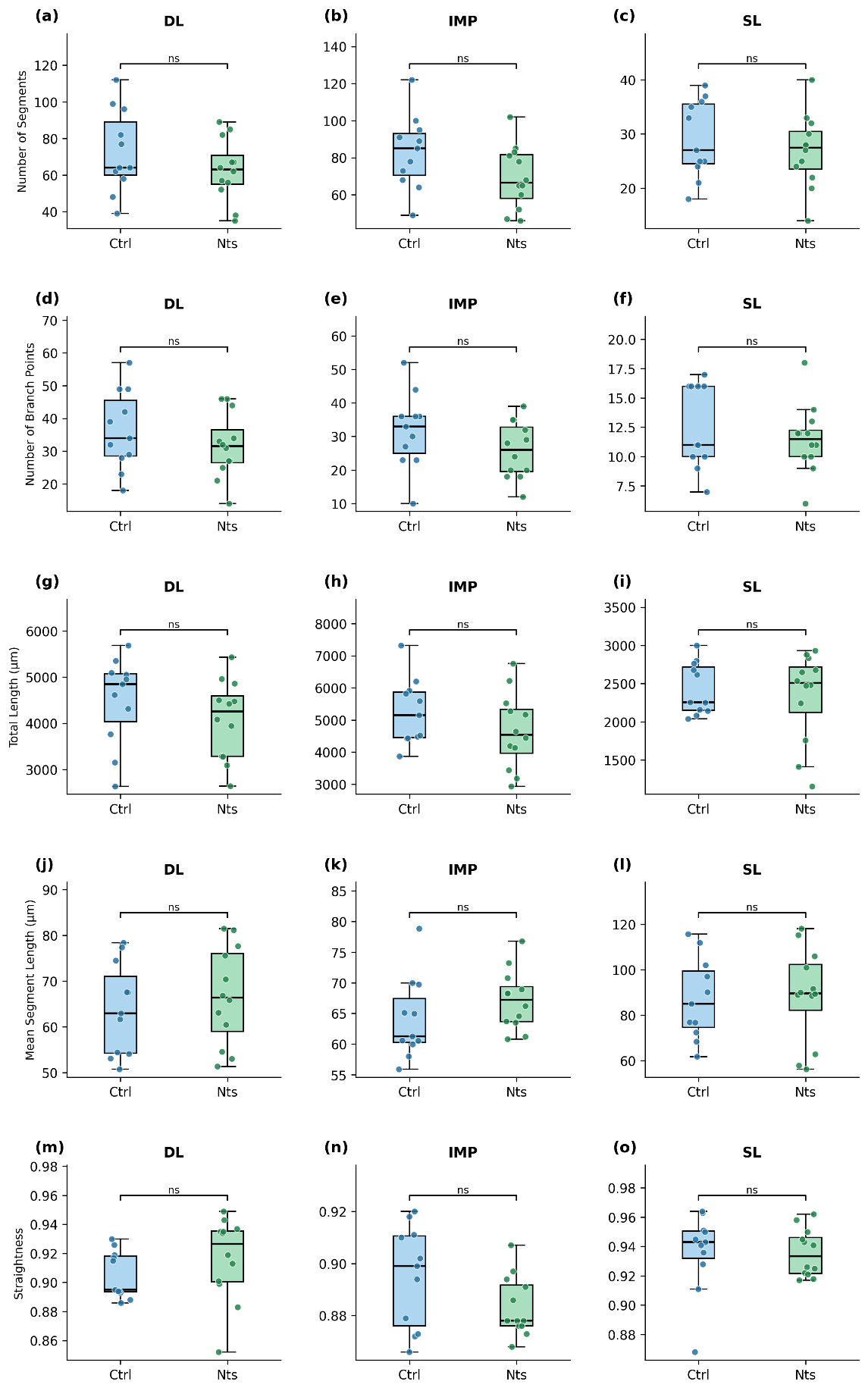

**Supplementary Figure S4. Additional plexus level vascular morphometric analyses in Piezo2^Nts^ retinas.** Vascular metrics were quantified separately for the deep layer (DL; panels a, d, g, j, m), intermediate layer plexus (IMP; panels b, e, h, k, n), and superficial layer (SL; panels c, f, i, l, o). (a–c) Total number of vessel segments. (d–f) Total number of branch points. (g–i) Total vessel length (sum of all segment lengths). (j–l) Mean segment length. (m–o) Vessel tortuosity (vessel length / end-to-end distance; higher values indicate greater curvature). Across all three plexus layers, no significant differences in tortuosity were observed between Control and Piezo2^Nts^ retinas.

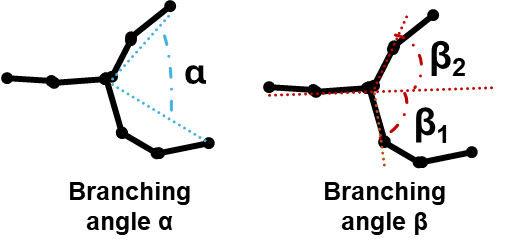

**Supplementary Figure S5. Branching angle definitions and hierarchical branch level classification.** Schematic illustration of vascular bifurcation geometry. α represents the angle between the two daughter vessel segments extending from the bifurcation point to their distal endpoints. β_1_ and β_2_ capture the angular deviation between the parent vessel’s extended centerline direction and each daughter vessel’s initial segment, measured at the nearest vertices from the bifurcation point.

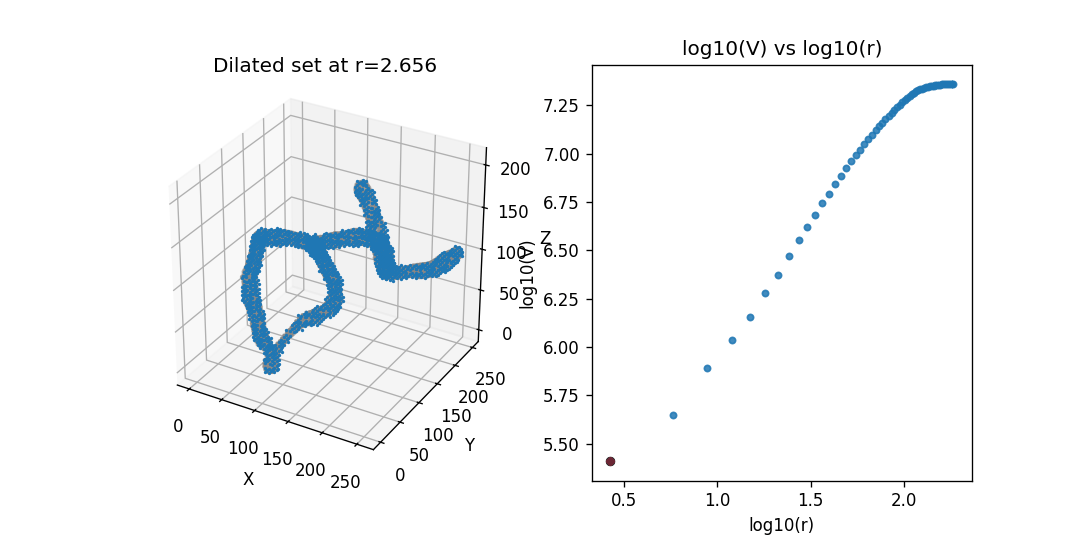

**Supplementary Figure S6. Demonstration of the Bouligand–Minkowski morphological dilation method for 3D fractal dimension computation.** Left: Progressive morphological dilation of a representative 3D vascular structure with spheres of increasing radius *r*. As *r* increases, the dilated set (blue) expands, and the influence volume *V*(*r*) grows accordingly. Right: Corresponding log_10_*V*(*r*) versus log_10_(*r*) plot. Each data point represents one dilation step. The fractal dimension is derived from the negative slope of the linear regression within the scale-invariant interval (Equation 1 in Methods). This visualization illustrates how the Bouligand–Minkowski method captures the space-filling complexity of 3D biological structures through systematic morphological analysis.

### Supplementary Video

#### Supplementary Video S1

**3D layer-resolved vascular quantification of control retina**

File: Video-S1-Ctrl_Layers Quantification.mp4

**Supplementary Video S1 |** Layer-resolved 3D vascular quantification workflow demonstrated on a representative control retina. The video illustrates the sequential stages of retinal vascular segmentation and layer stratification: full-thickness 3D reconstruction, layer stratification into deep (DL), middle (ML), and superficial (SL) plexuses, and visualization of layer-specific vascular architecture. The animation demonstrates how 3D imaging enables depth-resolved visualization of the trilaminar vascular organization that is not accessible through 2D en-face projections, highlighting the spatial relationships between plexuses that form the basis for inter-plexus connectivity and parallelism analyses described in the main text.

#### Supplementary Video S2

**3D rendered retina of col4a1 mice model control group**

File: Video-S2-Movie_Col4a1_Control.mp4

#### Supplementary Video S3

**3D rendered retina of CRA from col4a1 mouse model**

File: Video-S3-Movie_CRA.mp4

### Supplementary Methods

#### S.M.1 Inter-plexus vessel geometry: detailed computational procedures

**Orientation angle (θ) computation:**

For each penetrating vessel connecting two plexus layers, the orientation angle θ was computed as follows:

1. Identify the vessel’s fusion points with the upper and lower plexus layers.

2. Compute the straight-line vector **V** connecting these two fusion points.

3. Define the reference axis **N** as the axis orthogonal to the plexus planes (approximated by the superficial layer GAM-fitted surface normal at the nearest grid point).

4. Compute θ = arccos(|**V** · **N**| / (|**V**| · |**N**|)), yielding values from 0° (perfectly vertical/orthogonal) to 90° (parallel to plexus planes).

**Excursion ratio (ER) computation:**

1. For each penetrating vessel, trace the complete vessel path through all intermediate vertices from the upper to the lower fusion point.

2. Compute the vessel path length *L* as the sum of Euclidean distances between consecutive vertices along the vessel centerline.

3. Compute the orthogonal inter-plexus distance *D* as the vertical (z-axis) distance between the two plexus surfaces at the vessel’s x,y position.

4. Calculate ER = *L* / *D*, where ER = 1.0 indicates a perfectly direct (vertical) route and ER > 1 indicates increasing tortuosity or obliquity.

5. For statistical analysis and visualization, log_10_(ER) was used to normalize the distribution.

**Connection type classification:**

Inter-plexus connections were classified based on the originating and terminating plexus layers. For each penetrating vessel, the layers at both fusion points were identified through the manual layer stratification performed during segmentation. Connections were categorized as DL–ML (deep-to-middle), SL–ML (superficial-to-middle), or DL–SL (deep-to-superficial, bypassing the middle layer). Connection density was computed as the number of connections of each type divided by the analyzed area in the en-face plane.

#### S.M.2 Convex hull and alpha shape computation

Volume normalization was performed using two complementary approaches:

Convex hull volumes were computed from the 3D vascular point cloud (all vertex coordinates) using the Quickhull algorithm implemented in scipy.spatial.ConvexHull. The convex hull provides a standardized measure of the tissue volume occupied by the vascular network, enabling density calculations that account for variable ROI geometries.

Alpha shape analysis was additionally performed using trimesh and pyvista for cases requiring tighter boundary estimation. The alpha parameter was empirically determined for each dataset to minimize inclusion of non-vascularized tissue while maintaining a connected boundary.

#### S.M.3 Retina3D Vascular Analyzer pipeline user workflow

The Retina3D Vascular Analyzer pipeline is designed to be accessible to researchers without specialized computational training. The typical workflow proceeds as follows:

**Step 1:** Export validated 3D vascular reconstructions from Imaris as VRML (.wrl) model files and CSV statistics files containing vertex coordinates, edge connectivity, segment lengths, and diameters.

**Step 2:** Rename exported files according to standardized naming conventions. The pipeline provides in-software guidance and documentation for the required naming format: [SampleID]-[LayerInfo]_[Suffix].csv/wrl.

**Step 3:** Launch the pipeline through the Tkinter graphical interface, specify the input directory, and select the desired analysis modules.

**Step 4:** The pipeline automatically executes sequential processing stages: geometry preprocessing (convex hull and alpha shape computation), fractal dimension analysis (Bouligand–Minkowski method), and statistical analysis (whole-structure, layer-resolved, and inter-layer connectivity).

**Step 5:** Results are presented through interactive R Shiny dashboards for real-time exploration, with automated batch processing available for reproducible HTML report generation.
